# Gain-of-function genetic screen of the kinome reveals BRSK2 as an inhibitor of the NRF2 transcription factor

**DOI:** 10.1101/832279

**Authors:** Tigist Y Tamir, Brittany M Bowman, Megan J Agajanian, Dennis Goldfarb, Travis P Schrank, Trent Stohrer, Andrew E Hale, Priscila F Siesser, Seth J Weir, Ryan M Murphy, Kyle M LaPak, Bernard E Weissman, Nathaniel J Moorman, M. Ben Major

## Abstract

NFE2L2/NRF2 is a transcription factor and master regulator of cellular antioxidant response. Aberrantly high NRF2-dependent transcription is recurrent in human cancer, and conversely NRF2 protein levels as well as activity is diminished with age and in neurodegenerative disorders. Though NRF2 activating drugs are clinically beneficial, NRF2 inhibitors do not yet exist. Here we used a gain-of-function genetic screen of the kinome to identify new druggable regulators of NRF2 signaling. We found that the understudied protein kinase Brain Specific Kinase 2 (BRSK2) and the related BRSK1 kinases suppress NRF2-dependent transcription and NRF2 protein levels in an activity-dependent manner. Integrated phosphoproteomics and RNAseq studies revealed that BRSK2 drives AMPK activation and suppresses mTOR signaling. As a result, BRSK2 kinase activation suppressed ribosome-RNA complexes, global protein synthesis, and NRF2 protein levels. Collectively, our data establish the catalytically active BRSK2 kinase as a negative regulator of NRF2 via the AMPK/mTOR signaling. This signaling axis may prove useful for therapeutically targeting NRF2 in human diseases.

**Summary Statement:** BRSK2 suppresses NRF2 signaling by inhibiting protein synthesis through mTOR downregulation.

## Introduction

The transcription factor <u>n</u>uclear factor erythroid 2-<u>r</u>elated <u>f</u>actor <u>2</u> (NFE2L2, hereafter referred to as NRF2) is central to the cellular response to oxidative and electrophilic stress (Itoh et al., 2010; Suzuki and Yamamoto, 2015). When active, NRF2 provides strong cytoprotective functions by upregulating expression of: 1) xenobiotic metabolism enzymes, 2) phase II detoxification enzymes, 3) drug efflux pumps, and 4) the thioredoxin and glutathione antioxidant systems (Itoh et al., 2010; Kensler et al., 2007). Under homeostatic conditions, the E3 ubiquitin ligase adaptor <u>k</u>elch like <u>E</u>CH-<u>a</u>ssociated <u>p</u>rotein <u>1</u> (KEAP1) binds ETGE and DLG motifs within NRF2 to promote NRF2 ubiquitylation and proteasomal degradation (Cullinan et al., 2004; Itoh et al., 1999; Tong et al., 2006). Electrophilic attack of reactive cysteines within KEAP1 during oxidative stress suppresses KEAP1-dependent ubiquitylation/degradation of NRF2, resulting in NRF2 stabilization, nuclear translocation, and transcriptional activation of genes containing <u>A</u>ntioxidant <u>R</u>esponse <u>E</u>lements (AREs)(Baird et al., 2013; Ichikawa et al., 2009; Kensler et al., 2007; Suzuki et al., 2019). Though KEAP1/CUL3 are prominent NRF2 regulators in most tissues and disease models, data describing additional mechanisms are still emerging.

As a key regulator of metabolism and redox-balance, it is not surprising that alterations in NRF2 contribute to numerous human diseases. NRF2 expression and transcriptional activity declines with age and is decreased in several neurodegenerative diseases (Cuadrado et al., 2018; Tsakiri et al., 2019; Zhang et al., 2015). Conversely, NRF2 transcriptional activity is constitutively active in many cancers, including those of the lung, head/neck, kidney, liver, and bladder cancer (Menegon et al., 2016; Rojo de la Vega et al., 2018). Multiple mechanisms promote NRF2 activation in cancer: activating mutations in NRF2, inactivating mutations in KEAP1, NRF2 copy number amplifications, the over-expression of KEAP1 binding proteins which competitively displace NRF2, and various post-translational modifications to KEAP1 and NRF2(Cloer et al., 2019; Rojo de la Vega et al., 2018). High NRF2 activity promotes tumor growth, metastasis, and chemo-/radiation-resistance (Homma et al., 2009; Tao et al., 2014). Patients with mutations in KEAP1 or NRF2 leading to NRF2 stabilization have poor survival and are comparatively resistant to many commonly used cytotoxic cancer therapies(Homma et al., 2009; Satoh et al., 2013; Tao et al., 2014). While there are numerous small molecule activators of NRF2 signaling, there are no approved NRF2 inhibitors (Cuadrado et al., 2018).

Here, we performed a gain-of-function (GOF) screen of human kinases to identify new regulators of NRF2-dependent transcription. We focused on kinases given their exceptional druggability and because a rigorous, comprehensive annotation of the kinome for NRF2 activity is lacking. Our previous phosphoproteomic analysis of KEAP1 and NRF2 shows both proteins are phosphorylated at multiple sites. The majority of these phosphorylation events are not linked to specific kinases and are of unknown functional importance (Tamir et al., 2016). That said, recent studies have revealed a few kinases that influence NRF2 protein stability, subcellular localization and transcriptional activity. NRF2 is directly phosphorylated by GSK3β, resulting in NRF2 ubiquitylation by βTrCP and subsequent proteasomal degradation (Chowdhry et al., 2012; Cuadrado, 2015). PKC and AMPK mediated phosphorylation of NRF2, at S40 and S550, respectively, leads to increased NRF2 stability and signaling (Huang, 2002; Joo et al., 2016). NRF2 is reported to be a substrate of several MAPKs (e.g. JNK, p38, ERK1/2, ASK1, and TAK1), however the functional relevance of MAPK-directed phosphorylation is uncertain (Naidu et al., 2009; Shen et al., 2004; Sun et al., 2009). Activation of the PERK kinase during the unfolded protein response (UPR) leads to increased NRF2 nuclear accumulation and cell survival (Cullinan et al., 2003; Del Vecchio et al., 2014). The casein kinase 2 and TAK1 kinases phosphorylate NRF2 to induce NRF2 nuclear localization (Apopa et al., 2008; Shen et al., 2004). There is also evidence that activation of the PI3K/AKT/PKB pathway and PIM kinase signaling induces NRF2-activity and cellular protection (Lim et al., 2008; Nakaso et al., 2003; Warfel et al., 2016). Phosphorylation of KEAP1 or its interacting partners also induces NRF2 stability and signaling. For example, a recent study reports that phosphorylation of KEAP1 by MST1/2 on T51/S53/S55/S80 reduces NRF2 ubiquitylation (Wang et al., 2019). Additionally, proteins containing ETGE-like motifs, such as p62/SQSTM1, bind to KEAP1 and stabilize NRF2 upon phosphorylation by upstream kinases (e.g. mTORC1, TAK1) (Hashimoto et al., 2016; Ichimura et al., 2013; Lau et al., 2010). Overall, these findings suggest a wide array of kinase-directed signaling inputs for NRF2.

Our GOF kinome screen revealed that Brain Specific Kinase 2 and 1 (BRSK2/1, also known as SAD-A and SAD-B, respectively) as negative regulators of NRF2. The BRSKs are understudied members of the AMPK-related family of kinases(Bright et al., 2008). Both BRSK2/1 contain an N-terminal kinase domain, followed by a Ubiquitin associated domain (UBA), a Proline rich region (PRR), and a kinase associated domain (KA1) with an auto-inhibitory sequence (AIS) at the C-terminus (Wang et al., 2018; Wu et al., 2015). These kinases are known to function downstream of Liver Kinase B1 (LKB1) signaling, but are also activated by PAK1, CAMKII, and PKA (Bright et al., 2008; Lizcano et al., 2004; Nie et al., 2012). In various model organisms and mammals, BRSK2 and BRSK1 are expressed the most in the brain. BRSK2 is also expressed in pancreas; BRSK1 is expressed in gonads as well as endocrine tissues (Uhlen et al., 2015; Uhlen et al., 2010). In *C. elegans*, the *BRSK2/1* homologue *SAD-1* is required for neuron polarization and synaptic vesicle transport (Kim et al., 2010; Morrison et al., 2018). In mouse neurons, Brsk2/1 promote formation of vesicles, neurite differentiation, and synapse formation (Kishi, 2005; Lilley et al., 2014). In pancreatic islets, BRSK2 promotes insulin secretion in response to glucose stimulation (Chen et al., 2012; Nie et al., 2018; Nie et al., 2012). To date BRSK2/1 have been implicated in positive regulation of cell cycle progression, ER associated protein degradation (ERAD), neuronal polarity, autophagy, and insulin signaling (Li et al., 2012; Muller et al., 2010; Wang et al., 2012; Wang et al., 2013). Specifically, BRSK2 kinase activity is induced by starvation to inhibit mTOR and promote autophagy, similarly to AMPK, and new evidence suggests BRSK2 leads to activation of PI3K/AKT signaling (Bakula et al., 2017; Chen et al., 2012; Saiyin, 2017). While BRSK2 is not yet directly linked to NRF2 signaling, several studies show that BRSK2 is protective during ER stress (Wang et al., 2012; Wang et al., 2013). Here we identified BRSK2 as a novel repressor of NRF2 signaling, and mechanistically establish that BRSK2 activates AMPK, suppresses mTOR and decreases protein translation.

## Results

### Kinome gain of function screen identifies regulators of NRF2 activity

To identify kinase regulators of NRF2 signaling, we employed a gain-of-function arrayed screen where 385 kinases and kinase associated proteins were over-expressed in HEK293T cells. NRF2 transcriptional activity was quantified with the hQR41-Firefly luciferase reporter normalized to constitutively expressed *Renilla* luciferase. The hQR41 reporter contains a NRF2-responsive fragment of the human *NQO1* promoter, which is a consensus NRF2 target gene across species and tissue types. Over-expression of the positive controls NRF2 and DPP3 activated hQR41-luciferase, whereas KEAP1 expression suppressed the reporter activity (Hast et al., 2013). Kinases that activated hQR41 include MAP3K8, MOS, MAP3K7/TAK1, and MAP2K6; while HIPK4, BRD3, and BRSK2 were identified as repressors of NRF2-mediated transcription (Fig. 1A). Hits from this primary screen were validated in focused reporter assays across multiple cell lines, including HEK293Ts, MEFs and H2228 lung adenocarcinoma cells (Fig. 1B-E). Several of the validated kinases are confirmed in recent reports. MAP3K7/TAK1 activates NRF2 via phosphorylation of p62/SQSTM to drive degradation of KEAP1 (Hashimoto et al., 2016). Though BRD3 has not been directly tested, other members of the BRD family of proteins negatively regulate NRF2-mediated transcription (Hussong et al., 2014; Michaeloudes et al., 2014). Similarly, the HIPK4-related protein HIPK2 is a direct transcriptional target of NRF2 and a positive regulator of NRF2 cytoprotection (Torrente et al., 2017). Because of its novelty within the NRF2 pathway, we sought to better understand BRSK2 and its role in NRF2 signaling.

**Figure 1.**
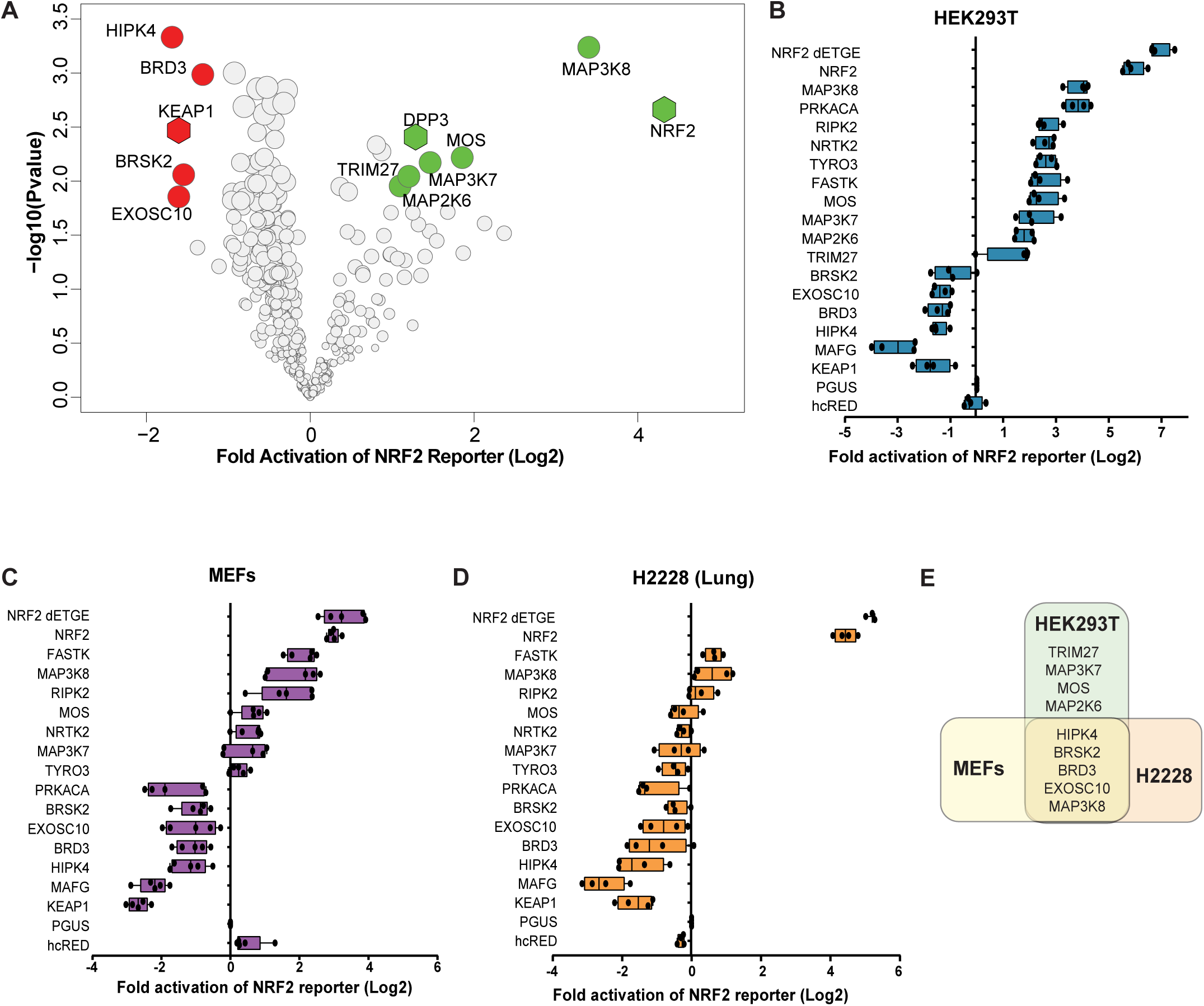
A gain-of-function screen identifies kinases that regulate NRF2-dependent transcription. **(A)**. Volcano plot representation of kinase over-expression screen of NRF2-driven hQR41-firefly luciferase reporter in HEK293T cells. The experiment was performed in biological triplicate and 6 technical replicates. NRF2 transcriptional activity was determined by measuring levels of hQR41-firefly luciferase normalized to TK-*Renilla* luciferase. Data are plotted relative to GFP over-expression control. Significance was measured with Student T-test including correction for multiple comparison using Benjamini & Hochberg method. Colored circles indicate candidate kinases that passed false discovery rate of less than 10%. NRF2, KEAP1 and DPP3 serve as positive controls. **(B-D)**. hQR41 reporter validation study for the indicated kinase in HEK293T cells, MEFs and H2228 cells, respectively. **(E)** Venn diagram of validated kinase hits in HEK293T, H2228 and MEFs.

### BRSK2 and BRSK1 repress NRF2 signaling

Redox regulation of cysteines within KEAP1 controls KEAP1-catalyzed NRF2 ubiquitylation and degradation. Through reactivity with cysteine 151 in KEAP1, the triterpenoid of oleanolic acid (OA) derivative CDDO-methyl (CDDO-me) potently suppresses NRF2 degradation (Suzuki et al., 2019). We first tested whether BRSK2 over-expression suppressed CDDO-me driven NRF2 activation, as quantified by hQR41-luciferase expression in HEK293T cells. We compared murine Brsk2 and two human BRSK2 splice variants, where BRSK2-Isoform4 is missing 20 amino acids compared to BRSK2-Isoform3. KEAP1 and Musculoapoeneuorotic Factor G (MAFG) served as positive controls for NRF2 inhibitors. Compared to the negative controls (hcRED and Glucuronidase Beta (pGUS)), all BRSK2 variants suppressed NRF2 transcriptional activity under both vehicle and CDDO-me treated conditions (Fig. 2A). Similarly, murine Brsk2 blocked NRF2-transcriptional activation. To confirm BRSK2 as an inhibitor of endogenous NRF2, we quantified endogenous NRF2 target gene expression by qRT-PCR. Following over-expression of controls or BRSK2 variants in HEK293T cells, we measured changes in NRF2 target genes *HMOX1, GCLM*, and *SLC7A11* normalized to *GAPDH*. BRSK2 variants downregulated NRF2 targets similar to MAFG in both vehicle and CDDO-me treated conditions (Fig. 2B-D). Since key regulation of NRF2 is post-translational, we tested whether BRSK2 over-expression impacted steady state NRF2 protein levels. Compared to controls, BRSK2 over-expression decreased NRF2 protein levels in HEK293T cells (Fig. 2E, compare lanes 5–7 with 1 and 2). To further explore a role for KEAP1 in mediating BRSK2-suppression of NRF2, we tested if BRSK2 over-expression could block a constitutively active mutant of NRF2 that does not bind KEAP1 (NRF2 ΔETGE). BRSK2 repressed both wild type NRF2 and NRF2 ΔETGE (Fig. 2F).

**Figure 2.**
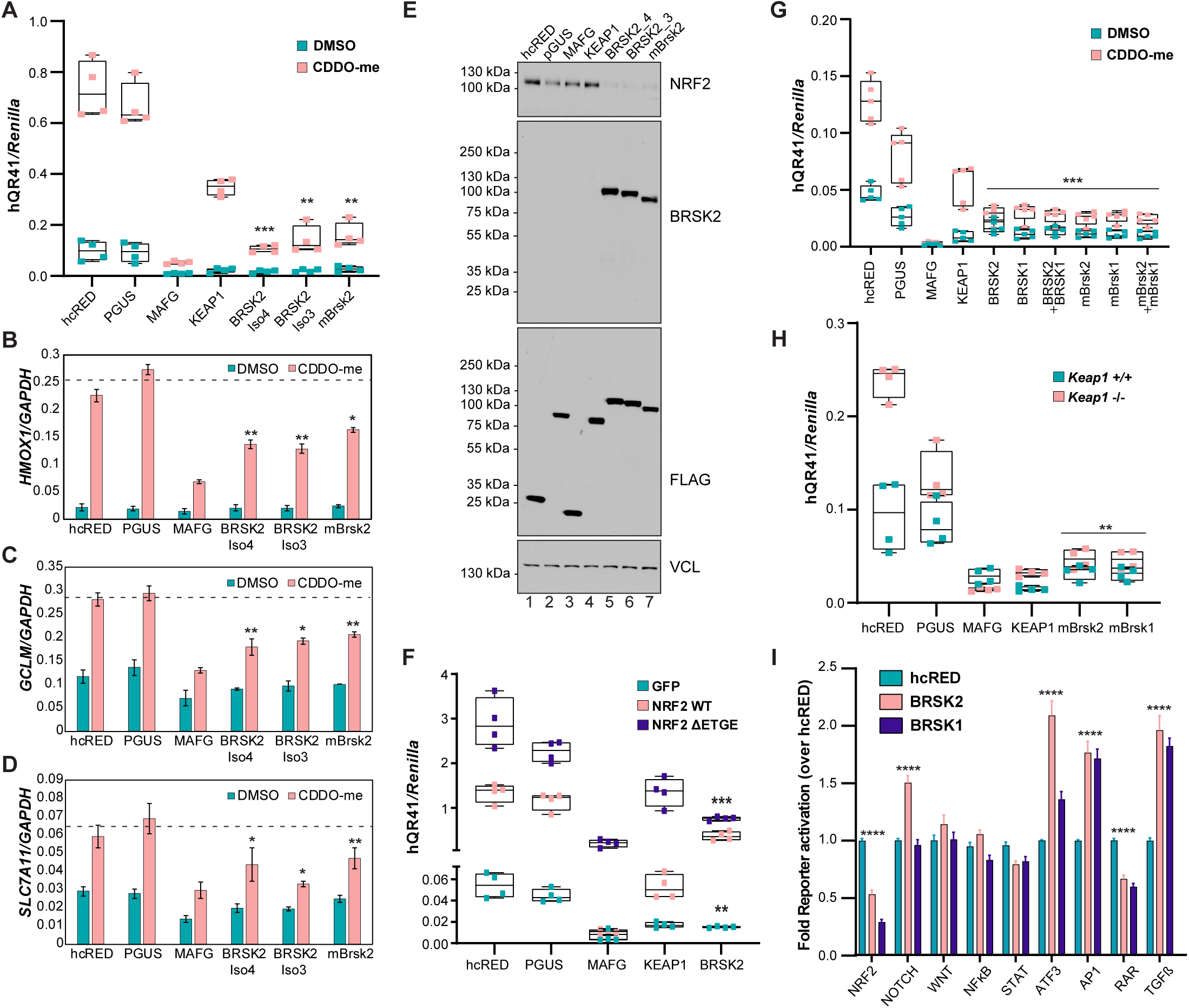
BRSK2 and BRSK1 inhibit NRF2-mediated transcription. **(A)** hQR41-luciferase assay in HEK293T cells following expression of control (hcRED, PGUS), positive control (MAFG, KEAP1) or BRSK2 splice variants (BRSK2iso4, BRISK2 iso3) or mouse Brsk2. Cell were treated with vehicle control or CDDO-me (100nM, 12hrs). **(B-D)** BRSK2 over-expression inhibits endogenous NRF2 target genes (*HMOX1, GCLM*, and *SLC7A11*) induction in HEK293T cells (representative of N = 3). (**E)** Western blot of HEK293T cells 24h after transfection with the indicated expression plasmid. Cells were treated with CDDO-me 12h before lysis. **(F)** hQR41-luciferase reporter assay in HEK293T cells expressing the indicated plasmid combination for 24h. **(G)** hQR41-luciferase reporter assay in HEK293T cells expressing the indicated plasmid. Cells were treated with CDDO-me 12h before lysis. **(H)** hQR41 reporter assay in wild type MEFs and KEAP1 -/- MEFs expressing the indicated plasmids. **(I)** HEK293T cells were transiently transfected with the indicated firefly transcriptional reporter, *Renilla* luciferase, hcRED, BRSK2 or BRSK1. Two-tailed Student’s t-test was performed comparing BRSK2 or BRSK1 to hcRED control, and statistical significance was assigned as follows: * p < 0.05, ** p < 0.01, *** p < 0.001, **** p < 0.0001.

BRSK1 is a paralog to BRSK2, sharing ∼68% amino acid sequence similarity and having similar signaling functions. Leveraging the hQR41 reporter assay in HEK293T cells, we over-expressed human or mouse BRSK1, BRSK2 or both. Like BRSK2, BRSK1 expression suppressed NRF2-dependent transcriptional activation (Fig. 2G). We also performed the hQR41 reporter assay in *Keap1* -/- MEFs, which express high levels of NRF2 (Fig. 2H). Again, both BRSK1 and BRSK2 inhibited NRF2-driven transcription, suggesting that the mechanism of suppression was independent of KEAP1-mediated ubiquitylation. Finally, we evaluated the impact of BRSK1 and BRSK2 on a panel of pathway specific transcriptional reporters in HEK293T cells (Fig. 2I). BRSK1 and BRSK2 suppressed NRF2 and retinoic acid receptor (RAR) and activated the Activator Protein-1 (AP1), Activating Transcription Factor 3 (ATF3), and Transforming Growth Factor β (TGFβ) reporters. Neither BRSK family member regulated the WNT/β-catenin reporter, NFκB reporter or STAT reporter. As such, BRSK1 and BRSK2 have redundant and conserved functions in NRF2 suppression, and do not impact all signaling pathways equally.

Finally, we evaluated the effect of BRSK2 silencing on NRF2 protein expression and NRF2-driven transcription. HEK293T cells were engineered to express dead KRAB-dCas9 nuclease before stable introduction of 4 scrambled control sgRNAs or 5 independent BRSK2-specific sgRNAs. W.blot analysis of the resulting cell lines confirmed efficient sgRNA-mediated CRISPRi silencing of BRSK2 protein and no effect on NRF2 protein levels (Fig. S1A). NRF2 hQR41 reporter assays in these cells did not show a BRSK2-silencing phenotype on NRF2 activity. Since *BRSK2* RNA levels are highest in brain and pancreas, we evaluated BRSK2 expression in pancreatic cancer cell lines (Fig. S1C). Two cell lines, PANC1 and MIA PaCa-2, expressed the most BRSK2, of which we used MIA PaCa-2 for CRISPRi silencing. Like HEK293T cells, MIA PaCa-2 cells deficient for BRSK2 expressed comparable levels of NRF2 as control cells (Fig. S1D). These data suggest that under homeostatic conditions, endogenous levels of BRSK2 does not control NRF2 activity. Further experiments in *BRSK2* null background as opposed to silenced background are needed.

### BRSK2 kinase function is required to suppress NRF2 signaling

We next determined whether BRSK2-mediated inhibition of NRF2 required BRSK2 kinase activity. We mutated K48 and D141 in the kinase domain of BRSK2; these residues are required for ATP binding and substrate phosphorylation (Lizcano et al., 2004). We also mutated T174, which is phosphorylated by LKB1 to activate members of the AMPK kinase family (Fig. 3A) (Lizcano et al., 2004). Compared to controls, over-expression of wild type BRSK2 decreased NRF2-dependent hQR41 luciferase expression whereas kinase dead variants had no significant affect (Fig. 3B). The kinase dependency of BRSK2 was further confirmed using qRT-PCR of endogenous NRF2 target gene *HMOX1*, which was repressed by wild type BRSK2, but not by the kinase dead mutants (Fig. 3C). qRT-PCR for the *NRF2* transcript showed that expression of neither wild type nor kinase dead BRSK2 mutants affected NRF2 mRNA levels (Fig. 3D). Finally, we evaluated NRF2 protein levels in HEK293T cells over-expressing BRSK2 or the kinase dead mutants. As expected, NRF2 protein levels were decreased by over-expression of wild type, but not kinase dead BRSK2 (Fig. 3E). Similarly, kinase dead mutants of BRSK1 did not impact NRF2 signaling (Fig. 3F).

**Figure 3.**
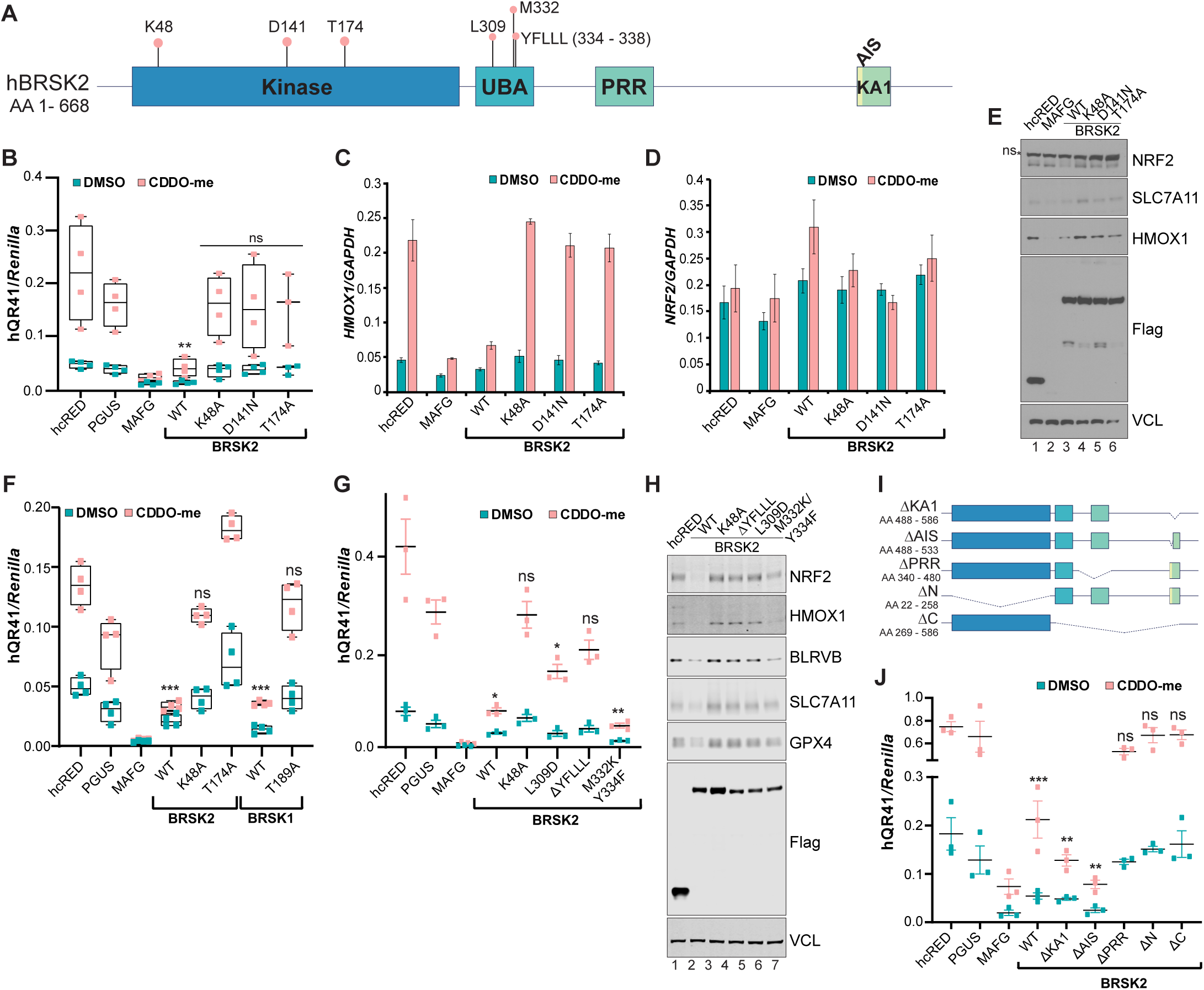
BRSK1 and BRSK2 kinase activity is required for suppression of NRF2-dependent transcription. **(A)** Cartoon schematic of BRSK2 protein showing residues and domains important for kinase activity. **(B)** hQR41 reporter assay in HEK293T cells expressing the indicated construct; cells were treated with vehicle or CDDO-me for 12h prior to luciferase quantitation. **(C**,**D)**. Quantitative RT-PCR for *HMOX1* and *NRF2* in HEK293T cells following over-expression of the indicated genes (representative of N = 3). **(E)** Western blot analysis of HEK293T cells expressing the indicated plasmids. Cells were treated with CDDO-me for 4h before lysis. ns = nonspecific. **(F)** hQR41 luciferase assay in HEK293T cells expressing the indicated proteins. **(G)** Western blot analysis of HEK293T cells expressing the indicated plasmids. Cells were treated with CDDO-me for 4h before lysis. **(H)** Cartoon schematic of BRSK2 deletions. **(I**,**J)** hQR41 reporter assay in HEK293T cells expressing the indicated construct; cells were treated with vehicle or CDDO-me for 12h prior to luciferase quantitation. Two-tailed Student’s t-test was performed comparing corresponding hcRED control with treatment groups and statistical significance was assigned: * p < 0.05, ** p < 0.01, *** p < 0.001, ns = not statistically significant.

Beyond kinase activity, structure-function studies of BRSK2 and the AMPK-like family of kinases have revealed numerous features of functional importance (Bright et al., 2009; Wu et al., 2015). We created a series of point mutations and truncations to determine which domains within BRSK2 are important for NRF2 suppression. Binding of the regulatory UBA and AIS region of the KA1 domain to the catalytic fold of the kinase domain is thought to hold BRSK2 in an inactive state until phosphorylated by upstream regulators, bound by interaction partners or recruited to the plasma membrane (Wu et al., 2015). As such, mutations in residues of the UBA domain and loss of the AIS or KA1 domain results in an active BRSK2 kinase. We created and expressed mutant proteins in HEK293T cells followed by hQR41 reporter quantitation and NRF2 W.blot analysis (Fig. 3G-J). The L309D and ΔYFLLL mutations within the UBA domain did not suppress the hQR41 reporter and NRF2 or its target genes in W.blot analysis (Fig.3G and 3H, respectively). Based on previous reports on the role of the BRSK2 UBA domain, it is likely that loss of YFLLL motif increases auto-inhibition by the AIS/KA1 region (Wu et al., 2015). Mutations of UBA residues that contact the kinase domain, M332K and Y334F, repressed NRF2 activity. We next deleted the kinase domain (ΔN), C-terminus (ΔC), PRR (ΔPRR), AIS (ΔAIS), or KA1 (ΔKA1) domain and evaluated their effect on NRF2-mediated transcription via reporter assays (Fig. 3I). Compared to controls, BRSK2 ΔKA1 and ΔAIS inhibited NRF2 activity to similar or more than BRSK2 WT, while loss of either kinase, C-terminal, or PRR regions abolished regulation of NRF2 signaling (Fig. 3J).

### BRSK2 does not repress NRF2 via the ubiquitin proteasome system (UPS)

NRF2 is rapidly stabilized by electrophilic compounds that react with cysteines in KEAP1. NRF2 protein levels also accumulate within minutes of chemical inactivation of the ubiquitin proteasome system (UPS). We tested whether BRSK2 expression would suppress NRF2 stabilized by KEAP1 reactive electrophiles or inhibitors of the UPS. First, we treated HEK293T cells with vehicle, Sulforaphane, or tBHQ for 6 hours after 24 hours of BRSK2 expression. Like CDDO-me, these compounds modify cysteine residues on KEAP1 to stabilize NRF2 (Suzuki et al., 2019). Cells over-expressing BRSK2 significantly downregulated NRF2 protein levels compared to the corresponding control (Fig. 4A, quantification below). Second, we tested the proteasomal inhibitors MG132 and Bortezomib as well as the CUL3 neddylation inhibitor MLN4924. NRF2 protein levels were decreased by BRSK2 in all treatment groups compared to the control, although to varying degrees (Fig. 4B, quantification below). Lastly, we asked whether BRSK2-mediated downregulation of NRF2 involved the autophagy pathway. To evaluate this, we treated BRSK2 overexpressing HEK293T cells with vehicle or Bafilomycin A (BafA1) for 12 hours. Compared to vehicle, BafA1 treatment did not significantly affect NRF2 in either control or BRSK2 over-expressing conditions (Fig. 4C, quantified below). LC3B conversion confirmed efficacy of BafA1 treatment. These data suggest that BRSK2-mediated downregulation of NRF2 is not via the UPS or autophagy.

**Figure 4.**
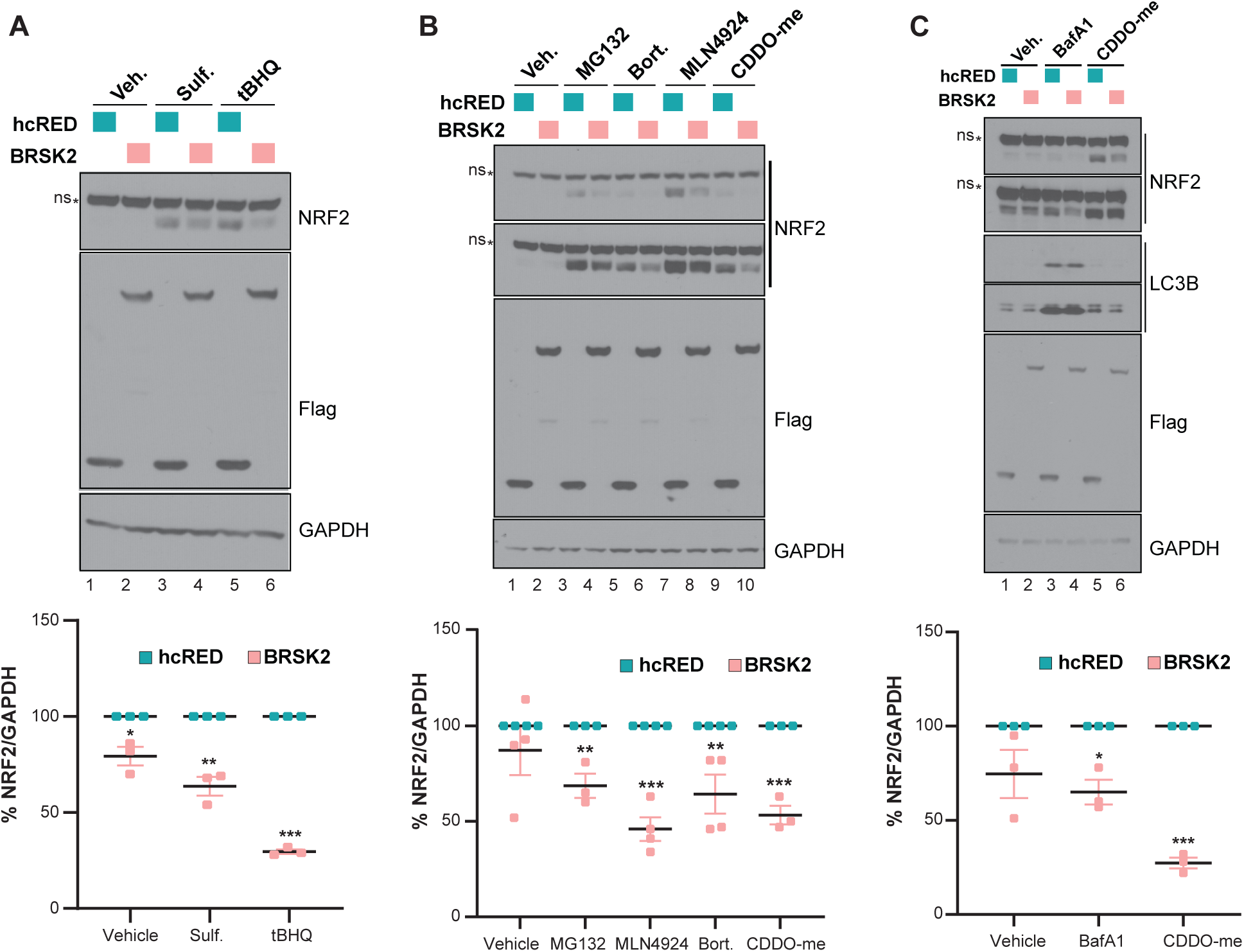
BRSK2-mediated NRF2 downregulation is independent of the NRF2 ubiquitylation and degradation. **(A)** HEK293T cells were transfected with control hcRED or BRSK2 for 24h before treatment with vehicle (veh) or Sulforaphane (Sulf., 2μM) or tert-Butylhydroquinone (tBHQ, 50μM) for 4h. Quantitation of biological triplicate experiments is shown in the lower panel, normalized to GAPDH. **(B, C)** HEK293T cells were transfected with control hcRED or BRSK2 for 24h before treatment with MG132 [10μM], Bortezomab (Bort., [40nM]), MLN4924 [2.5μM], Bafilomycin A1 (BafA1, [200nM]). All treatments were for 4h, except for BafA1 which was 12h. Two-tailed Student’s t-test was performed comparing hcRED with BRSK2, and statistical significance was assigned as follows: * p < 0.05, ** p < 0.01, *** p < 0.001.

### RNAseq and phosphoproteomic characterization BRSK2 and BRSK1 expression

Unbiased comprehensive screening and molecular annotation has improved significantly over the past decade, with newly empowered informatics that distill the resulting large datasets into pathways and biological processes. To better understand BRSK2/1 function in cells, and to possibly reveal how BRSK2 suppresses NRF2, we performed RNAseq and global quantitative phosphoproteomic analysis on HEK293T cells expressing BRSK2 and BRSK1 as compared to mock transfected or hcRED expression. Hierarchical clustering analysis revealed genes altered by BRSK2/1 over-expression, where genes that passed FDR (FDR < 5%) and fold change (FC ≥ 2) cutoff are highlighted (Fig. 5A and B). Compared to control, we observed 723 and 879 differentially expressed genes that pass FDR < 5% for BRSK2 and BRSK1, respectively. Based on fold change, differential gene expression due to BRSK2 expression was more robust than BRSK1. Gene set enrichment analysis (GSEA) using the Hallmark and Oncogenic gene sets in MSigDB revealed statistically significantly altered signaling pathways (Table S5 – S8). Genes associated with mTOR signaling were robustly downregulated in BRSK2/1 expressing cells (Fig. 5C). GSEA for several NRF2 gene signatures revealed downregulation by BRSK2 (Fig. S2, Table S9). Close examination by pathway analysis of the downregulated genes revealed enrichment for those involved in pyruvate, glutathione, and amino acid metabolism. Interestingly, several of the genes are known players in ferroptosis, a non-canonical and iron dependent cell death pathway (Table S9).

**Figure 5.**
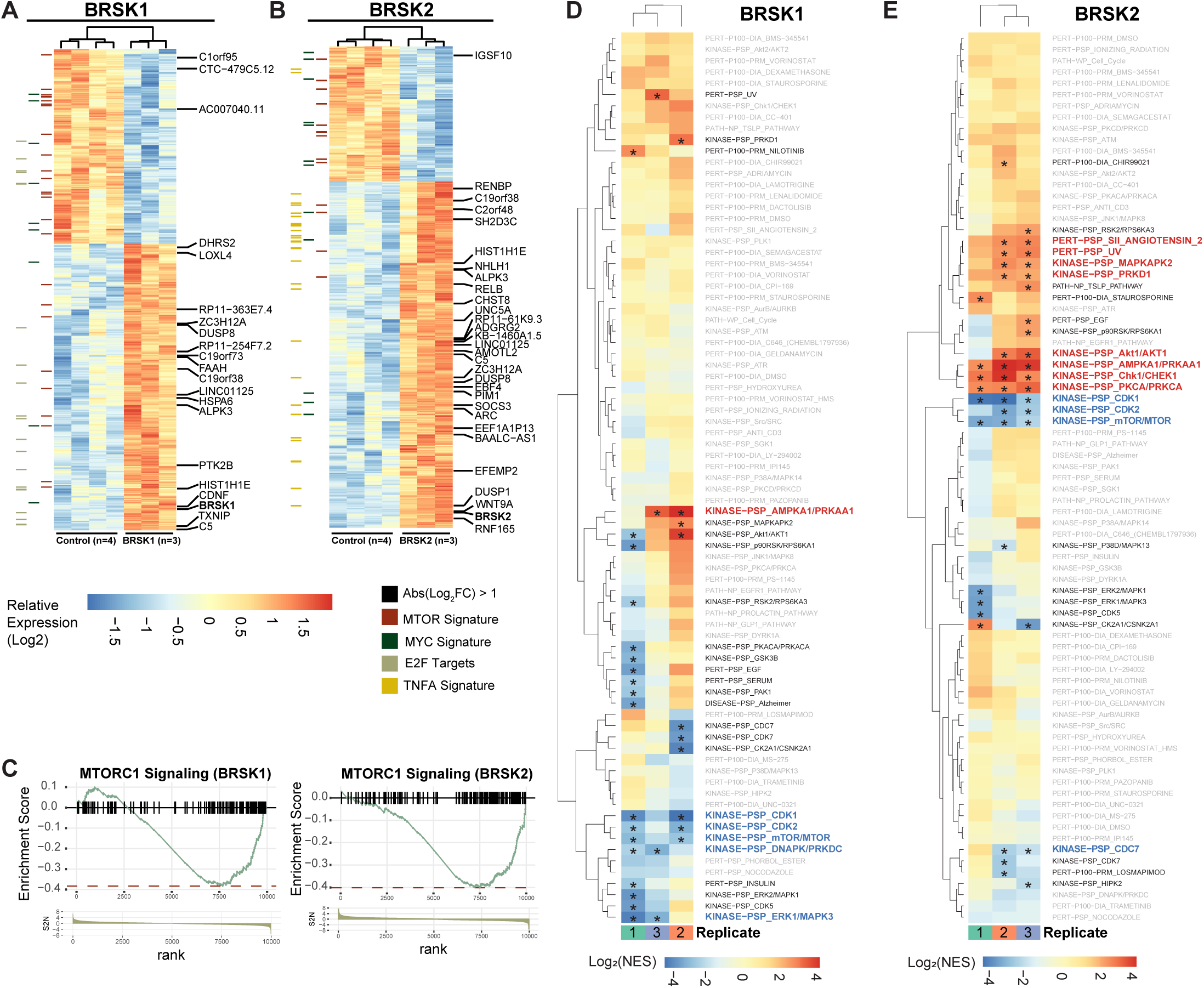
Genomics and proteomics reveals activation of AMPK and suppression of MTOR following BRSK1/2 expression. **(A, B)** Hierarchical clustering of RNA sequencing analysis of HEK293T cells transfected with mock, hcRED control or BRSK2 or BRSK1 for 24h. **(C)** Differentially expressed genes with FDR < 5% were analyzed by Gene Set Enrichment Analysis (GSEA) using the Hallmark and Oncogenic gene sets from MSigDB. Enrichment Score (ES) plots represent genes commonly associated with MTORC1 signaling were decreased by BRSK2/1 overexpression based on GSEA. **(D, E)** Quantitative TMT phosphoproteomic analysis of HEK293T cells expressing hcRED, BRSK1, or BRKS2 for 24h. TMT ratios from biological triplicate samples were analyzed for enrichment of phosphosites associated with known signaling pathways using the PTMSigDB pipeline (* FDR < 10%).

Independently of the RNAseq analyses, we performed tandem mass tags (TMT)-based quantitative phosphoproteomics on BRSK2/1 expressing cells where we used hcRED as control. Biological triplicate samples were analyzed, revealing ∼10,000 phosphosites in ∼8,400 phospho-peptides. Following the RNAseq trend, BRSK2 impacted phospho-proteome more robustly than did BRSK1. Compared to control, at FDR < 5%, BRSK2 over-expression induced 307 differentially phosphorylated peptides compared to 189 observed in BRSK1 over-expression (Table S10). We leveraged PTMSigDB and enrichment analysis to map the observed phospho-peptide changes to annotated signaling pathways (Fig. 5D and E) (Krug et al., 2019). BRSK2/1 positively regulated AMPK and AKT signaling, while negatively impacting the mTOR pathway. BRSK2 also suppressed signaling through the CDK1, CDK2, and CDC7 pathways (Table S12).

To confirm activation of AMPK signaling, BRSK2 was expressed in HEK293T cells before W.blot analysis for phosphorylation of AMPK substrates (LxRxx(pS/pT)). We expressed either wild type (WT), kinase active (ΔKA1 or ΔAIS), or kinase dead (K48A or D141N) BRSK2 for 24 hours. Over-expression of wild type and kinase active BRSK2 (lane 2, 5, & 6) upregulated pS/T AMPK motif compared to control and kinase dead BRSK2 (lane 1, 3, & 4) (Fig 6A). We then measured changes in mTOR substrate phosphorylation by evaluating p-S6K T389, p-4EBP1 T37/46, and p-4EBP1 S65 in cells expressing hcRED control, BRSK2, kinase dead BRSK2 (K48A), or BRSK1. BRSK2/1 expression decreased phosphorylation of both S6K and 4EBP1 (Fig. 6B). Total protein levels of 4EBP1 increased following BRSK2/1 expression. Together, these data suggest that BRSK2/1 downregulates mTOR signaling while upregulating AMPK substrate phosphorylation.

**Figure 6.**
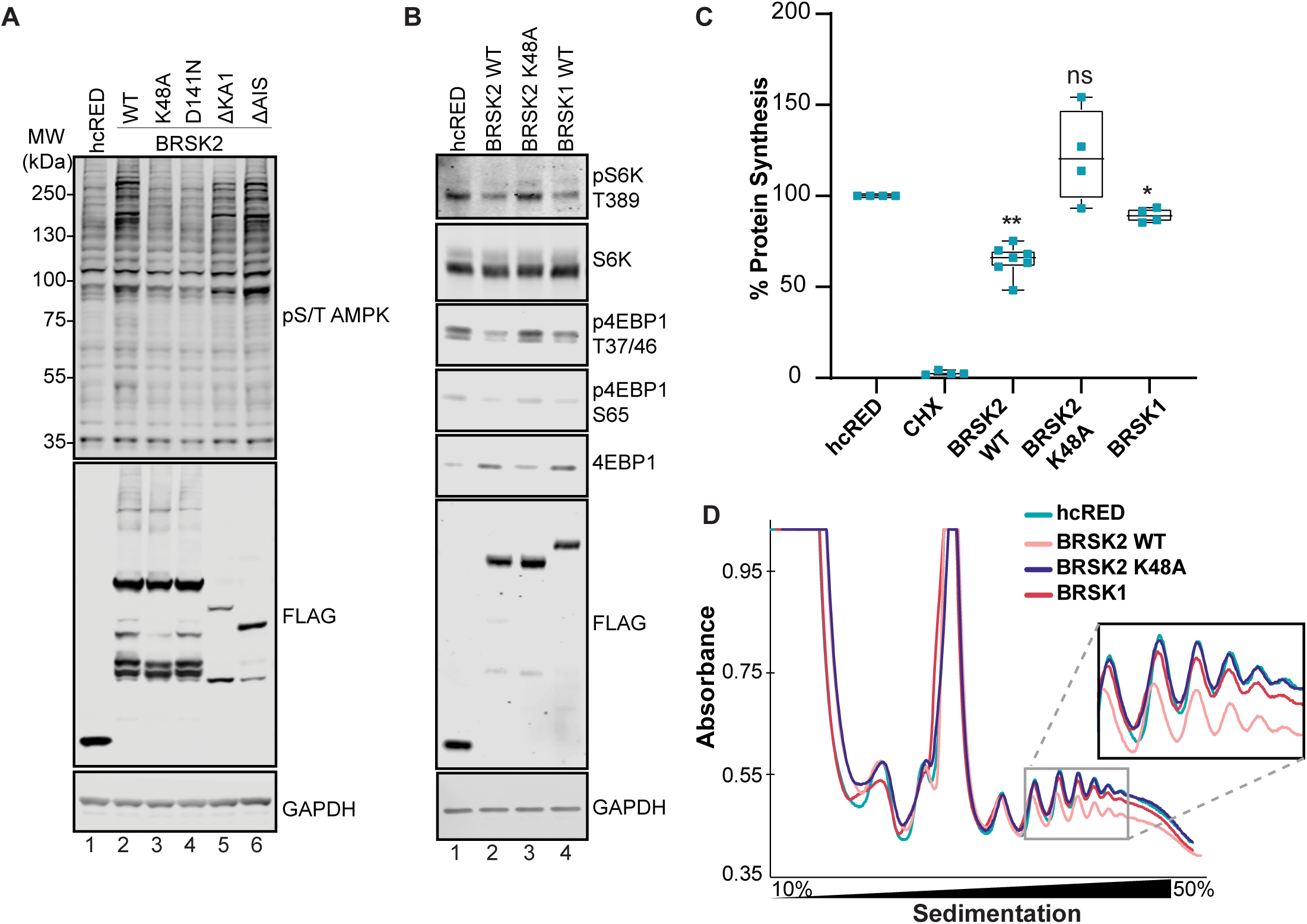
BRSK2 suppresses protein translation. **(A)** Western blot analysis AMPK consensus phosphorylation sites in HEK293T cells transfected with hcRED, BRSK2, or the indicated BRSK2 mutants (representative of N = 3). **(B)** Western blot analysis of BRSK1 and BRSK2 expression in HEK293T cells for MTOR targets S6K (T389) and 4EBP1 (T37/46 and S65) which increases 4EBP1 stability (representative of N = 3). **(C)** Global protein translation assay performed using ^35^S-Metionine labeling in cells expressing hcRed (control), BRSK2, BRSK2 K48A, or BRSK1. Two-tailed Student’s t-test was performed comparing hcRED to other conditions, and statistical significance was assigned as follows: * p < 0.05, ** p < 0.01, *** p < 0.001. **(D)** Polyribosome fraction of HEK293T cells overexpressing hcRED control, BRSK2, BRSK2 K48A, or BRSK1 performed on 50% – 10% sucrose gradient. Experiments represent biological triplicates, unless indicated otherwise.

Finally, because BRSK2/1 over-expression suppressed mTOR, we quantified the rate of protein translation after BRSK2/1 expression (Fig. 6C). HEK293T cells expressing BRSK2/1 or control hcRED were pulsed with ^35^S-Methionine before lysis and quantitation of nascent polypeptides, where Cyclohexamide (CHX) served as a negative control. Compared to control and kinase dead BRSK2, expression of wild type BRSK2 and BRSK1 downregulated protein translation by 40% and 10%, respectively (Fig. 6C). To further confirm decreased translation, we measured ribosome binding to mRNA in cells expressing BRSK2/1. Cells transfected with the indicated construct were fractionated in a 10% – 50% sucrose gradient followed by absorbance measurement for polyribosome tracing. Compared to control and kinase dead BRSK2, over-expression of wild type BRSK2 decreased heavy polyribosome formation on mRNA (Fig. 6D).

## Discussion

The NRF2 transcription factor is central to a growing number of human pathologies. The transcriptional program it governs enables life in the presence of oxygen, electrophiles, and environmental stressors. As such, aberrant activation or suppression of NRF2 contributes to and causes a number of human diseases, including cancer, inflammation, diabetes, and neurodegeneration. Though its relevance and centrality to human health is well-established, we have yet to realize the full potential of NRF2-directed therapeutics. In this study, we focused on kinase regulators of NRF2 because the kinome is exceptionally druggable and the mechanistic impact of NRF2 phosphorylation remains elusive. We discovered that, in an activity-dependent manner, the BRSK2 kinase suppresses NRF2 signaling. We show that BRSK2 induces AMPK activity and inhibits mTOR, resulting in decreased ribosome loading on mRNAs which downregulates protein synthesis.

Beyond revealing the BRSK kinases as indirect suppressors of NRF2 translation, this work has several implications. Signaling through the mTOR pathway has been reported to both promote and inhibit NRF2 in a context dependent manner (Aramburu et al., 2014). Our data argue that mTOR suppression by BRSK2, irrespective of cell type or duration, results in NRF2 protein loss. Specifically, BRSK2 over-expression suppresses NRF2 protein at basal and electrophilic induced conditions. Though NRF2 suppression was greatest after 24 hours of BRSK2 expression, through an unknown mechanism, BRSK2 over-expression for greater than 36 hours resulted in cell death (not shown). While we connect BRSK2 to NRF2 through activation of AMPK and suppression of mTOR, it is possible that additional mechanisms contribute to the phenotype. For example, it remains to be determined if BRSK2/1 bind or directly phosphorylate KEAP1 or NRF2. Our immunoprecipitation and mass spectrometry experiments for KEAP1, NRF2, BRSK1 and BRSK2 did not reveal a co-complex, but false-negatives are common in pull-down mass spectrometry experiments, particularly for kinase-substrate interactions.

The small molecule NRF2 inhibitors halofuginone and brusatol offer further support for a requisite role of protein synthesis as a key point of NRF2 regulation (Harder et al., 2017; Tsuchida et al., 2017). Both halofuginone and brusatol are noted as potent NRF2 inhibitors that sensitize cells to chemotherapy. Halofuginone inhibits prolyl-tRNA synthetase while brusatol acts on peptidyl transferase to suppress protein translation and ultimately decrease proteins with short half-life like NRF2, a phenotype observed with over-expression of BRSK2 (Harder et al., 2017; Tsuchida et al., 2017). This also highlights the underlying role of stress signaling in protein synthesis. Under moderate oxidative stress, mTOR regulated 5’ Cap-dependent translation is inhibited, but can be bypassed via Internal Ribosomal Entry Site (IRES)-mediated translation found in the 5’ untranslated region (5’UTR) (Komar and Hatzoglou, 2011). However, under severe stress translation is inhibited via localization of ribosome-RNA complexes to stress granules to protect nescient polypeptides and translation complex from damage. While we have not evaluated changes in NRF2 mRNA levels associated with polysomes, the NRF2 mRNA does contain an IRES-like region that promotes de novo translation under oxidative stress (Lee et al., 2017).

Conversely, NRF2 loss decreases protein translation rate in pancreatic cancer due to increased oxidation of proteins in the translational machinery as well as decreased 4EBP1 phosphorylation (DeNicola et al., 2012). Since BRSK2/1 decreases mTOR activity, possible mechanism of NRF2 depletion may be block of translation initiation/elongation under low oxidative stress where NRF2 mRNA is still dependent on 5’Cap-dependent translation.

Gain-of-function screening and extensive validation confirm BRSK2 as a suppressor of NRF2, yet it is unclear if BRSK2 loss impacts NRF2 or oxidative stress response signaling. In an acute experiment, we designed and tested a panel of short interfering RNAs against BRSK2, and observed inconsistent results despite greater than 95% BRSK2 silencing (not shown). Long-term suppression of BRSK2 expression with stable CRISPRi technology did not impact NRF2 protein levels. Since kinases are catalytic enzymes, it is likely that our loss-of-function approaches failed to eliminate enough BRSK2 activity to impact NRF2. Alternatively, it is possible that our experiments lacked the necessary context, for example the presence of an upstream activating signal. Further studies are needed in neuronal and pancreatic systems to evaluate BRSK loss and NRF2 signal transduction.

BRSK2 and BRSK1 impact cellular signaling pathways beyond NRF2, AMPK and mTOR, as gleaned from the RNAseq and phosphoproteomic experiments. BRSK2 expression activated CHEK1, PKC, and AKT1 among other kinase signaling pathways. In a focused screen of various engineered transcriptional reporters, we show that BRSK1 and BRSK2 induced AP1 and ATF3 stress response signaling, as well at TGFβ signaling. It is possible that some of these responses are coupled indirectly to the suppression of mTOR; further work is needed to determine this. These data provide a foundational resource to better interrogate signaling regulated by BRSK2/1.

## Material and Methods

### Cell Culture

All cell lines were obtained from American Type Culture Collection (ATCC) and were used within 10 passages after receipt from ATCC to ensure their identities. Cells were cultured in a humidified incubator at 37°C and 5% CO_2_. Cells were passaged with 0.05% Trypsin/0.53mM EDTA in Sodium Bicarbonate (Corning, 25-052-CI), and maintained in media supplemented with 10% fetal bovine serum (FBS) as follows: HEK293T, MIA PaCa2– Dulbecco’s Modified Eagle Medium (DMEM) (Corning, 10-013-CV); H2228– RPMI-1640 (Corning, 10-040-CV); and *Keap1* +*/*+ and *Keap1 -/-* Mouse Embryonic Fibroblasts (MEFs) generated as previously described (Cloer, 2018; Wakabayashi et al., 2003) – DMEM/Ham’s F-12 50/50 mix supplemented with Sodium Pyruvate and Non-essential amino acids (Corning, 10-092-CV).

### Generation of CRISPRi Cell lines

Lentivirus for KRAB-dCas9 was generated by using PsPax2 and PMD2G packaging vectors (Gilbert et al., 2014). HEK293T and MIA PaCa-2 cells were infected with KRAB-dCas9 lentivirus and monoclonal lines were generated via single cell sorting following 10µg/mL Blasticidin (GIBCO, A11139-03) selection for 5 passages. Each monoclonal line was cultured in 5µg/mL Blasticidin following sorting. Single guide RNA (sgRNA) vectors were generated by ligating oligonucleotides into AarI (ThermoFisher Scientific, ER1582) digested VDB783 vector (Table S1). Each sgRNA was lentivirally introduced to the above mentioned cell lines, and cells were cultured for 3 passages in 2.5µg/mL Puromycine (Corning, 61-385-RA) and 5µg/mL Blasticidin before further analysis. Stable cell lines were maintained in 1µg/mL Puromycine and 5µg/mL Blasticidin.

### Plasmids and Reagents

The human kinome ORF library in pDONOR223 was obtained from Addgene (Cambridge, MA, 1000000014) and cloned into a custom pHAGE-CMV-FLAG destination vector using Gateway cloning technology, as previously described (Agajanian et al., 2019). Luciferase reporter plasmids for WNT (BAR), NFκB, AP-1, ATF3, STAT, Retinoic acid (RAR), and TGFβ (SMAD) were cloned into transient expression vectors as previously done (Matthew P. Walker, 2015; Travis L. Biechele, 2008). The NOTCH (CSL) reporter was a gift from Raphael Kopan (Addgene, 41726) (Saxena et al., 2001).

All ORFs were cloned into pHAGE-CMV-FLAG via LR clonase (Thermo Fisher, 11791-020): pDONOR223.1 *BRSK2* (splice isoform 3, BRSK2_Iso3) and *Brsk2* were obtained from Harvard PlasmID repository (HsCD00297097 and MmCD00295042). BRSK2 Kinase dead (K48A, D141N, and T174A) and UBA domain (L309D, M332K/Y334F, and ΔYFLLL) mutants were generated using Q5 Hot Start Site Direct Mutagenesis kit (New England BioLabs, E0552S). BRSK2 domain deletion mutations ΔN (Δ kinase), ΔC (Δ C-terminal), ΔPRR (Δ proline-rich region), ΔKA1 (Δ kinase associated domain), and ΔAIS (Δ auto-inhibitory sequence) were generated via Phusion DNA polymerase (New England BioLabs, M0530S), where the PCR product was treated with DpnI (New England BioLabs, R0176S) then purified with Monarch PCR & DNA Purification Kit (New England BioLabs, T1030S) followed by T4 DNA Ligase reaction (New England BioLabs, M0202S). BRSK1 WT and T189A vectors were obtained from MRC PPU Reagents and Services (DU1236 and DU1242) in a pCMV5-HA backbone. Brsk1 was obtained from Origene (MR220008). They were then gateway converted into pDONOR223.1. Cloning primers are listed in Table S1.

### Kinome gain-of-function screen

HEK293T cells were plated in 384-well clear-bottom plates (Corning, 3764), and transfected with a cocktail of 20ng FLAG-Kinase, 4ng hQR41 (Moehlenkamp and Johnson, 1999) (a generous gift from Jeffery Johnson, University of Wisconsin), and 1ng HSV-thymidine kinase promoter driven *Renilla* luciferase (pRL-TK-*Renilla*, referred to as *Renilla*) (Promega, E2241) per well using Lipofectamine 2000 (Life technologies, 11668-019) in OptiMEM (Gibco, 31985-070). Each kinase was transfected in four technical replicates and biological triplicate using the Promega Dual-Glo Luciferase Assay System per the manufacturer’s protocol (Promega, E2940) (Agajanian et al., 2019). NRF2 transcriptional activity was determined by measuring levels of Firefly luciferase and was normalized to *Renilla* luciferase for well-to-well variability. Fold activation of the reporter was determined compared to GFP expressing cells, and false discovery rate (FDR) was calculated following adjustment via Benjamini & Hochberg correction.

### Western blot

All samples were lysed in RIPA (10% glycerol, 50mM Tris-HCL, 100mM NaCl, 2mM EDTA, 0.1% SDS, 1% Nonidet P-40, 0.2% Sodium Deoxycholate) supplemented with protease inhibitor cocktail (ThermoFisher Scientific, 78429), phosphatase inhibitor cocktail (ThermoFisher Scientific, 78426), NEM (Thermo Scientific, 23030), and Benzonase (Sigma, E1014). Lysis was done on ice for 30min and lysates were centrifuged at 4°C for 15min at 21,000xg. Following normalization of protein concentration via BCA (Pierce, 23225), samples were denatured in NuPAGE LDS buffer (Invitrogen, NP0007) with 1mM DTT. Samples were treated with the following small molecules to stabilize NRF2: Sigma-- Bardoxolone Methyl/CDDO-me (S8078), tert-Butal hydroquinone/tBHQ (112941), Sulforaphane (S6317); Calbiochem-- MG132 (474790), MLN4924 (505477001); Selleck Chem-- Bortezomib (PS-341). Antibodies are listed in Table S2.

### qRT-PCR and RNA sequencing analysis

RNA was collected using PureLink RNA Mini Kit (Invitrogen, 12183018A) per manufacturer instruction. The extracted RNA was then reverse transcribed to cDNA using iScript cDNA Synthesis Kit (BioRad, 170-8891), which was then used to perform quantitative RT-PCR (qRT-PCR) using SYBR Green (Applied Biosystems, 4385617) for the specified target genes (Table S3).

3µg of RNA was submitted to Novogene Corp. Ltd. (Sacramento, CA) for sequencing using Illumina HiSeq platform where reads were mapped to reference genome *Homo sapiens* (GRCh37/hg19). Alignments were done using STAR/HTSeq. Differential expression analysis was performed starting with gene level read count quantification provided by Novogene Corp. Preprocessing, normalization, and differential expression analysis were performed according to the analysis pipeline outlined by Law *et. al*. (Law et al., 2016). Marginally detected genes (<5 total read counts across samples) were filtered out as an initial preprocessing step, yielding a uni-modal distribution of read-counts per million by gene. Data normalization was performed by using the trimmed mean of M-values method as implemented in the calcNormFactors edgeR R-package function(Chen, 2014). Differential expression analysis was subsequently performed using the Limma R-packages functions voom and eBayes (Ritchie et al., 2015).

Gene set enrichment analysis (GSEA) was performed based on the above described pre-processed read counts which were converted to counts per million and log2 transformed (logCPM). From 13,000 genes the top 10,000 differentially expressed genes were used for GSEA analysis. Genes were ranked according to signal to noise ratio as defined by the Broad Institute GSEA software using the R-project fgsea package. Test gene sets (Hallmark and Oncogenic gene sets from Molecular Signatures Database (MSigDB, Broad Institute)) were downloaded from the MSIG data bank via the msigdbr R-project package (Liberzon et al., 2015; Subramanian, 2005). Raw data and processed datasets are available through GEO (GSE139135).

### Translation assay and polysome fractionation

Rate of protein translation was measured in HEK293T cells expressing controls or BRSK2/1 24 hours post transfection using radioactive methionine (^35^S-Met) labeling, and polysome fractionation was performed as previously described (Graves et al., 2019; Lenarcic et al., 2014).

### Phosphoproteomics sample processing and data analysis

Protein from HEK293Ts expressing control or BRSK2/1 (1.4mg) was precipitated in acetone overnight at −20°C. The sample was pelleted and re-suspended in 7M urea, reduced with 5mM DTT (dithiothreitol) and alkylated with 15mM CAA (chloroacetamide). The sample was adjusted with 50mM ABC (ammonium bicarbonate) such that the urea concentration is 1M or less. A standard tryptic digest was performed overnight at 37°C. Solid Phase Extraction (SPE) was then performed using C18 Prep Sep™ cartridges (Waters, WAT054960), followed by reconstitution in 0.5% TFA (trifluoroacetic acid). The SPE cartridge was washed with conditioning solution (90% methanol with 0.1% TFA), then equilibrated with 0.1% TFA. The sample was passed slowly (1drop/sec) through the equilibrated cartridge, then the cartridge was desalted with equilibration solution. The sample was then slowly eluted (1drop/sec) with an elution solution (50% ACN (acetonitrile)) with 0.1% TFA. The sample was then TMT labeled according to kit specifications (ThermoFisher Scientific, 90110), with the exception that labeling was performed for 6hrs instead of 1hr. Following labeling, another SPE was performed, as stated above. 10% of sample was saved for whole proteome input, 90% was phosphopeptide enriched using Titansphere Phos-TiO Kit (GL Sciences, 5010-21312). Before enrichment, samples were reconstituted in 100µL of Buffer B (75% ACN, 1% TFA, 20% lactic acid – solution B in the kit). The tip was conditioned by centrifugation with 100µL of Buffer A (80% ACN, 1% TFA), followed by conditioning with Buffer B (3000xg, 2min). The sample was then loaded onto the tip and centrifuged twice (1000xg, 5min). The tip was then washed with 50µL of Buffer B, followed by 2 washes with 50µL of Buffer A (1000xg, 2min). Phosphopeptides were eluted with 100µL of elution 1 (20% ACN, 5% NH4OH) then 100µL of elution 2 (20% ACN, 10% NH_4_OH) (1000xg, 5min). Following phosphopeptide enrichment, both whole proteome input and phosphopeptides were fractionated utilizing a High pH Reversed-Phase Peptide Fractionation kit (Pierce, 84868), per manufacturers specifications. A final clean-up step was performed using C18 Spin Columns (Pierce, 89870).

### Mass Spectrometry, Data Filtering, and Bioinformatics

Mass spectrometry analysis done as follows: peptides were separated via reverse-phase nano-HPLC using nanoACQUITY UPLC system (Waters Corporation). Peptides were trapped on a 2 cm column (Pepmap 100, 3μM particle size, 100 Å pore size; ThermoFisher Scientific, 164946), and separated on a 25cm EASYspray analytical column (75μM ID, 2.0μm C18 particle size, 100 Å pore size; ThermoFisher Scientific, ES802) at 45°C. The mobile phases were 0.1% formic acid in water (Buffer A) and 0.1% formic acid in ACN (Buffer B). A 180-minute gradient of 2-30% buffer B was used with a flow rate of 300nl/min. Mass spectral analysis was performed by an Orbitrap Fusion Lumos mass spectrometer (ThermoFisher Scientific). The ion source was operated at 2.4kV and the ion transfer tube was set to 275°C. Full MS scans (350-2000 m/z) were analyzed in the Orbitrap at a resolution of 120,000 and 4e5 AGC target. The MS2 spectra were collected using a 0.7 m/z isolation width and analyzed by the linear ion trap using 1e4 AGC target after HCD fragmentation at 30% collision energy with 50ms maximum injection time. The MS3 scans (100-500 m/z) were acquired in the Orbitrap at a resolution of 50,000 with a 1e5 AGC, 2 m/z MS2 isolation window, and at 105ms maximum injection time after HCD fragmentation with a normalized energy of 65%. Precursor ions were selected in 400-2000 m/z mass range with mass exclusion width of 5 – 18 m/z. Lock mass = 371.10124 m/z (Polysiloxane).

MaxQuant (1.6.6.0) search parameters: specific tryptic digestion, up to 2 missed cleavages, a static carbamidomethyl cysteine modification, variable protein N-term acetylation, and variable phospho(STY) as well as methionine oxidation using the human UniProtKB/Swiss-Prot sequence database (Downloaded Feb 1, 2017). MaxQuant data was deposited to PRIDE/Proteome Xchange (PXD015884) (Vizcaino et al., 2014). MaxQuant output files proteinGroups.txt and Phospho(STY) sites.txt were converted using in-house script into GCT format. This GCT file was then rewritten using Morpheus for compatibility with PTM-SEA analysis in R (Krug et al., 2019).

## Acknowledgements

We would like to thank Drs. Michael J Emanuele and Thomas R Bonacci for valuable scientific discussion. We thank Drs. Kirsten L Bryant and Channing J Der for providing expertise and resources to explore mTOR signaling as well as autophagy.

## Funding sources

TYT and MJA were funded by HHMI Gilliam Fellowship for Advanced Study and the Initiative for Maximizing Student Diversity Grant (R25-GM055336-16). Additionally, TYT was supported by NIH T32 Pre-doctoral Training Grants in Pharmacology (T32-GM007040-42). MJA was also funded via NIH/NCI NRSA (F31CA228289) and NCI F99/K00 (1F99CA245724-01). RMM was supported by NIH/NCI NRSA (1F31DE028749-01) and NIH/NIGMS T32 MiBio Training Program (5T32GM119999-03). BMB, TPS, and KML were funded by ITCMS T32 Training Grant (T32CA009156). BMB was also supported by NIH/NCI F32 (1F32CA225040-01). This research was supported by American Cancer Society grant to M. B. Major (RSG-14-1657068-01-TBE), NIH/NCI to M. B. Major and B. E. Weissman (RO1CA216051) and a NIH consortium grant on Illuminating the Druggable Genome (U24-DK116204-01).

## Author Contributions

This work was designed by MBM and TYT. Experiments were carried out by TYT, BMB, MJA, AEH, PSF, SJW, RMM, and KML. Bioinformatics analysis were performed by DG, TPS, TS, and TYT. Expertise and resources for evaluating protein translation biology were provided by NJM and AEH. Manuscript was written by TYT and MBM. Manuscript was reviewed and edited by BMB and BEW.

## Competing interests

Authors do not have competing interests to declare.

---

**Supplemental Figure S1.**
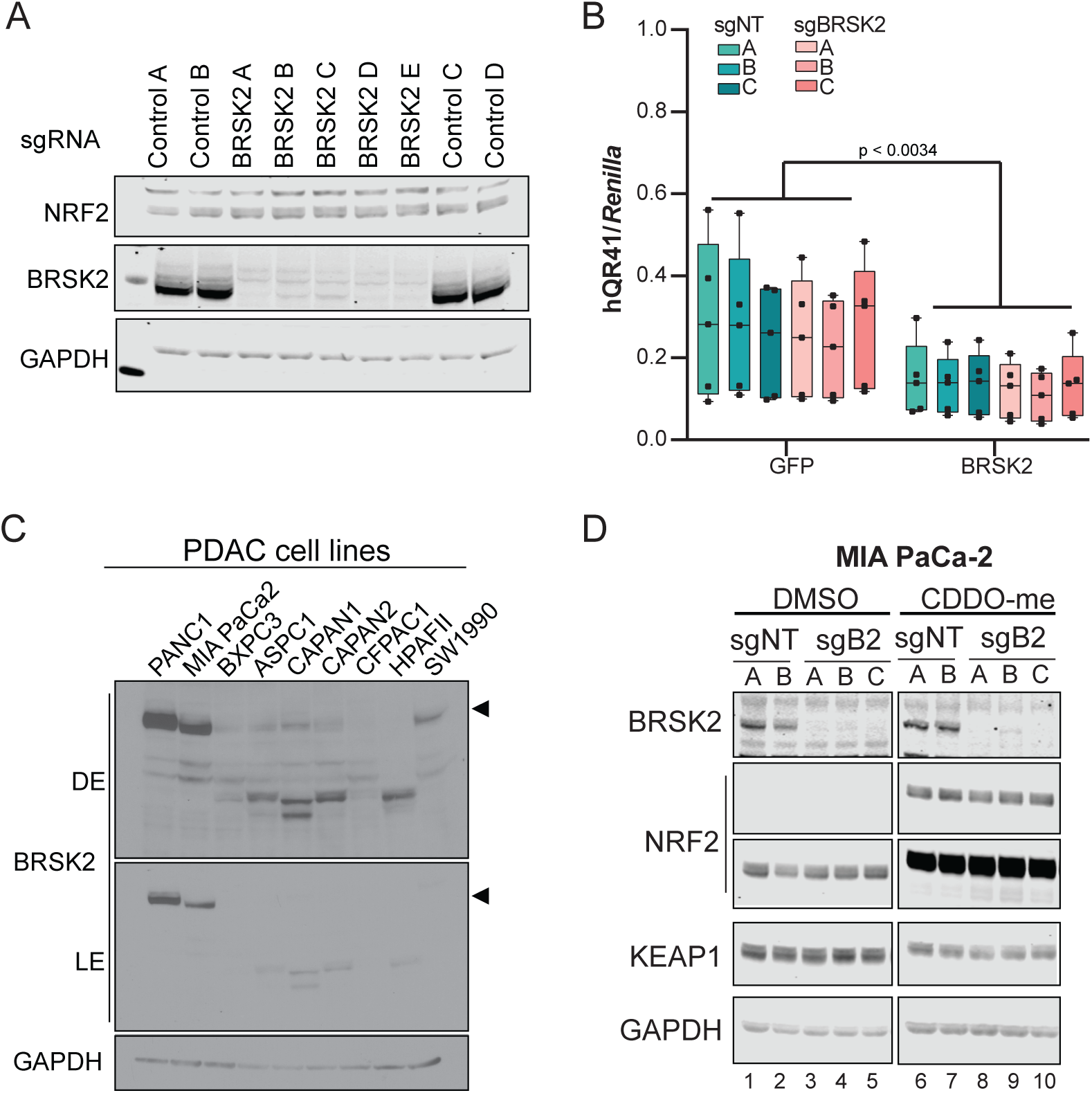
BRSK2 CRISPRi silencing does not alter NRF2 protein levels. **(A)** HEK293T cells stably expressing KRAB-dCas9 and 4 control or 5 BRSK2 sgRNAs. BRSK2 knockdown does not alter NRF2 protein levels. **(B)** hQR41 reporter assay in CRISPRi silenced cells with GFP or BRSK2 over-expression shows that BRSK2 knockdown does not affect NRF2 activity. **(C)** BRSK2 expression levels in pancreatic ductal adenocarcinoma (PDAC) cell lines. **(D)** CRISPRi silencing of BRSK2 in MIA PaCa2 cells treated with DMSO or CDDO-me for 4hrs. Neither NRF2 nor KEAP1 levels were altered by loss of BRSK2 protein.

**Supplemental Figure S2.**
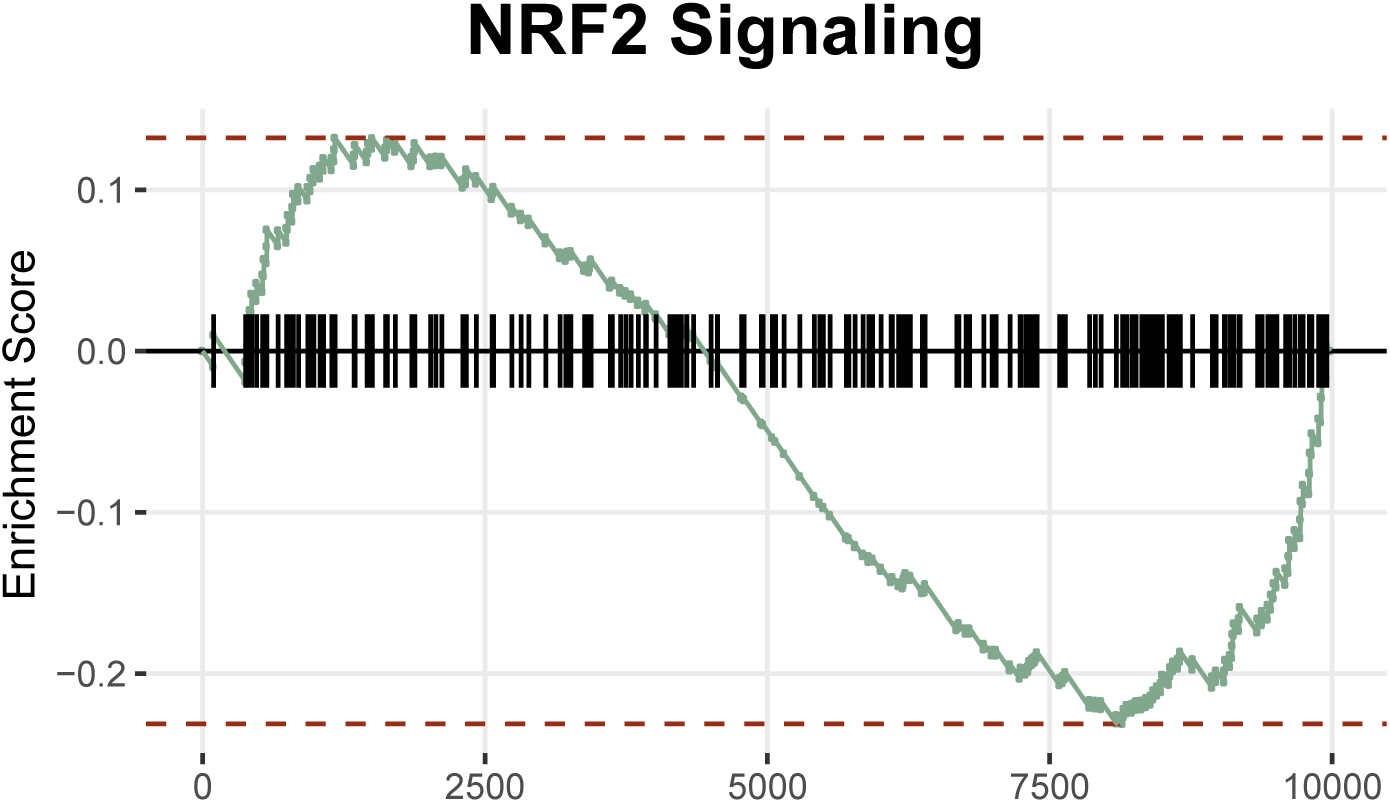
BRSK2 overexpression decreased NRF2 target gene signature.

